# Insect biomass of protected habitats under the impact of arable farming in Germany

**DOI:** 10.1101/2023.09.24.559203

**Authors:** Roland Mühlethaler, Sebastian Köthe, Thomas Hörren, Martin Sorg, Lisa Eichler, Gerlind U. C. Lehmann

## Abstract

Five years after the well-known study on insect biomass decline in nature protected habitats in Germany over three decades, the project DINA (Diversity of Insects in Nature protected Areas) has investigated the status of insects in 21 selected nature reserves across Germany in the years 2020 and 2021. We used the same methods and protocols for trapping and measuring the biomass of flying insects as in the mentioned study. Across two seasons, we accumulated a comprehensive data set of 1621 data points of two-week emptying intervals. The measured overall insect biomass remained at low levels and corresponds to the published latest figures from the years 2007–2016. There were no significant regional differences, but biomass was negatively correlated with agricultural production area within 2 km of nature reserves. Differences between the two consecutive years were very likely due to well-known natural fluctuations of insect populations, changes in agricultural cultivation and local weather events. The results show that protected habitats are essential for insects, but not sufficient in their function, and that further steps need to be taken for a better protection and sustainment of insects, which fulfil key functions in many ecosystems.

## Introduction

In the past few decades, alarming declines of insects have been observed in many places all over the world (Burner et al., 2021; Powney et al., 2019; Seibold et al., 2019). The study of Hallmann et al. (2017) illustrated the decline of insect biomass in nature protected areas (NPA) in Germany over 27 years, triggering major international impact. **Fehler! Textmarke nicht definiert.** The subsequent discussions made a significant contribution to raising the awareness of the general public and politicians about the problem of insect decline. Since then, several studies proved that insect biomass decline involves a loss of their diversity (Hallmann et al., 2021; Hausmann et al., 2022). In addition, fragmentation and decrease of insect populations as part of the observed insect decline is a major problem in many regions (Baur et al., 2020; Conrad et al., 2004).

NPA serve as legally binding areas for the special protection of nature and landscape in their entirety or in parts in order to enable the conservation, development or restoration of habitats, biotopes or communities of certain wild animal and plant species. Therefore, the protection of site-typical biodiversity should have unrestricted priority for all biotope types and types of land-use (including arable land biotopes). However, the German red list of threatened habitat types (Finck et al., 2017) shows that without exception all arable habitat types including a high degree of the wild arable plant communities are classified as “threatened with complete extinction”. A similarly dramatic situation is evident for the threat to wildlife in these arable habitats. With agricultural use as a major cause of insect decline (Sánchez-Bayo & Wyckhuys, 2019), NPA and special areas of conservation (SAC) of the European Natura 2000 network are becoming increasingly important for the conservation of biodiversity, including arable habitats (Eichler et al., 2022; Sorg et al., 2019).

The DINA project (Diversity of Insects in Nature protected Areas), a comprehensive interdisciplinary research project involving eight institutions (Lehmann et al., 2021), investigates the influence on insect populations in nature reserves as part of SAC and within the European Natura 2000 network (Brühl et al., 2021; Eichler et al., 2022; Köthe et al., 2023a; Lehmann et al., 2021; Swenson et al., 2022). Based on the original model to study the main drivers for the decline in insect biomass (Hallmann et al. 2017), we focused on the already identified covariates: the influence of arable land and the chemical inputs such as nitrogen and pesticides associated with intensive land use. In contrast to Hallmann et al. (2017), a meta-analysis across several individual studies and geographically limited regions, DINA has conducted a comprehensive inclusion at comparable sites (NPA) and investigated gradients through insect trap transects for the first time. Moreover, due to the comparability in methodology, the data can be collated with those of Hallmann et al. (2017). In addition to insect data, landscape elements and their spatial distribution pattern (Eichler et al., 2022), vegetation (Köthe et al., 2023b; Swenson et al., 2022), and pesticide residues were measured (Brühl et al., 2021) accompanied by exchange with local stakeholders for developing solutions for improved insect protection (Fickel et al., 2020; Köthe et al., 2023a; Lehmann et al., 2021).

Here we investigate the biomass data of two consecutive years 2020 and 2021 from 1621 Malaise trap samples in 21 NPA across Germany. The application of almost identical methods used by the Entomological Society Krefeld (Hallmann et al., 2017, 2021; Sorg et al., 2019; Ssymank et al., 2018) allows a comparison with the previous period from 1989-2017 and shows how the recent low insect biomass is distributed within high-quality NPA across Germany.

## Material and Methods

### Sampling sites and Malaise trap setup

Based on spatial analyses, landscape indicators were first evaluated, resulting in a pre-selection of sampling sites from a total of 8836 nature reserves in Germany (Lehmann et al., 2021). In addition to landscape indicators, the prerequisites for the sites were high-quality grassland-dominated habitat types of Natura 2000 (supplement Table S2) with adjacent agricultural areas and the cooperation of local authorities and landowners. Malaise traps have proven to be a reliable method for effective sampling of insects (Sorg et al., 2019). In 2020 and 2021, Malaise traps were set up in 21 nature reserves located within the borders of special areas of conservation (SAC) as part of the European Natura 2000 network across Germany to investigate insect communities from early May to late August with collection intervals of two weeks. Five Malaise traps were set at each site, creating a transect that started on arable land (Malaise trap 1 = MT1), continued along the boundary between agricultural land and protected area (MT2), and extended into the protected area (MT 3–5), with approximately 25 metres between each trap (Figure 1; see also Lehmann et al., 2021). At some sites, landowners withdrew their consent to place a Malaise trap on their fields, even during the current sampling period. At four sites in both years and at a further three in 2021, the first trap was therefore tipped at the boundary between agricultural land and protected area, corresponding to Malaise trap 2 (MT2a and MT2b in those cases). At one site (Brauselay, Rhineland-Palatinate), the first trap had to be completely removed in 2021 because the local situation did not allow relocation.

**FIGURE 1.**
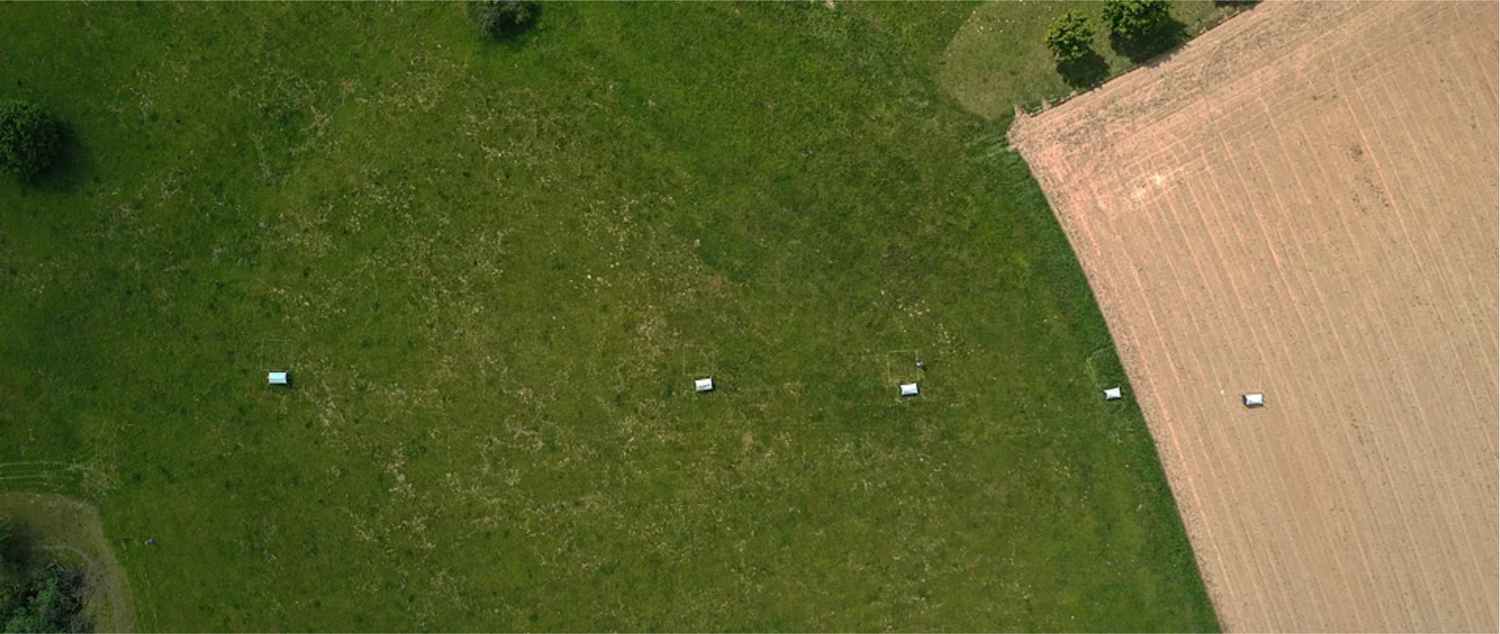
Aerial view of a Malaise trap transect starting with MT1 on arable land (right in picture), MT2 at the boarder of the protected habitat and on the left MT5 towards the centre of the nature reserve (copyright: Entomological Society Krefeld).

Comparability of data with past and future insect monitoring studies was ensured by using the standardised Malaise traps (type Townes, 1972) of the Entomological Society Krefeld (Sorg et al., 2019; Ssymank et al., 2018). These traps collect flying insects in 1000 ml polyethylene bottles, preserved in 96% ethanol from which biomass was determined (Hallmann et al., 2017; Ssymank et al., 2018). The ethanol containing the catches of insects and other arthropods is poured over a stainless-steel fine sieve (mesh size < 0.5 mm), which allowed unwanted by-catches such as snails (Pulmonata) to be sorted out. After the frequency of the ethanol drop exceeds approx. 10 seconds, the fresh biomass is measured on a fine balance (T500Y, G&G GmbH Neuss, lower measuring accuracy <0.1 g). Animals and ethyl alcohol are then returned to the original polyethylene bottles. For better comparison and because trapping intervals varied in some cases a few days, we standardized the wet biomasses as daily values.

### Climate data

Data on maximum temperature and daily precipitation were obtained from the German Weather Service (DWD). For each site, the closest weather station (distances ranging from 2.5 to 22.8 km from the Malaise trap site) was identified to collect information. As indicator for the overall climate conditions, we used the continentality index provided by the DWD (2021) for each study site.

### Statistical analysis

Statistical analysis was carried out using Rx64 4.0.1 (R Core Team, 2019). DINA is unique in applying transects of traps, including borders and adjacent arable land, in the sampling scheme. In order to compare our data with previous similar studies conducted in German NPA, we restricted several analyses to our traps located in protected habitats. For these analyses, the data of the three traps MT3–MT5 were pooled into a single mean. Generalized additive models (GAM) analysed the effects of maximum temperature, precipitation, continentality index, and agricultural production area within 2 km on daily biomass using the “mgcv” package fitted with the Restricted Maximum Likelihood (REML) method (Wood, 2011, 2017). Since the data for the continentality index and the agricultural production area within 2 km did not differ for MT3–5, and to avoid pseudoreplication in the modelling, the average daily biomass of traps per site was averaged for traps MT3 to MT5. The average maximum temperature and average precipitation were calculated accordingly.

Due to massive change in land use (e.g. newly established wild flower strips, flowering fallows) at the four sites Riedensee, Insel Koos, Bottendorfer Hügel and Schwellenburg in 2021 near the trap position where intensive arable farming was practiced in 2020, we performed two different GAM analysis per year. One with all 21 sampling sites and one excluding these four sites. For an analysis of local insect biomass dominance, we clustered the transect according to the placement of the traps into four classes: *arable land* (AL at MT1), *border* with traps directly at the boundary of the nature protected area (Bor at MT2), within the *nature protected areas* (NPA with MT3–MT5) and *indistinguishable* (Ind, at least two trap classes show similarly high values).

Four factors were tested as predictive variables of insect biomass in four generalized additive models (GAM), separated between years 2020 and 2021 and either including MT3-MT5 within NPA from all 21 localities or excluding the four sites with landscape change between years (supplement Table S4).

## Results

The mean daily insect biomass from a total of 1621 samples (829 from 2020 and 792 from 2021) varied from 0.95 (SE ± 0.47) to 3.62 (SE ± 1.98) grams per day (Supplementary Table S1). These biomasses varied considerably from year to year (Supplementary Figure S1), with insect biomass decreasing at 13 sampling sites and increasing at eight sites from 2020 to 2021 (Table 1, Supplementary Figure S2).

**TABLE 1.** Categorized mean maximum insect biomass of different Malaise traps along transects for 2020 and 2021: On *arable land* (AL), *border* between arable land and nature protected area (Bor), within *nature protected area* (NPA), *indistinguishable* (at least two trap classes show similarly high values) (Ind). Interannual unchanged dominance patterns in 12 out of 21 areas are indicated in blue.

| Sites | 2020 | 2021 | Difference in biomass 2020 /2021<br>(in %) |
| --- | --- | --- | --- |
| Lütjenholmer Heidedünen | Bor | Bor | –8.8 |
| Riedensee | NPA | AL | +51.4 |
| Insel Koos | NPA | AL | –2.7 |
| Geesower Hügel | Bor | Bor | +11.2 |
| Oderhänge Mallnow | AL | AL | +20.0 |
| Wisseler Dünen | NPA | Bor | –10.6 |
| Bislicher Insel | NPA | Ind | +4.7 |
| Gipskarstlandschaft Hainholz | Ind | Bor | –19.3 |
| Porphyrlandschaft bei Gimritz | Ind | Bor | –19.9 |
| Ziegenbuschhänge bei Oberau | NPA | NPA | –4.5 |
| Wipperdurchbruch | Ind | Ind | –1.0 |
| Bottendorfer Hügel | Bor | Ind | –7.1 |
| Schwellenburg | Bor | Bor | +30.1 |
| Hofberg | NPA | NPA | –22.8 |
| Koppelstein-Helmestäl | NPA | Ind | + 4.4 |
| Rheinhänge Dörscheider Heide | Bor | Ind | + 5.1 |
| Brauselay | NPA | NPA | -21.2 |
| Mittelberg | Ind | Ind | -42.4 |
| Ipf | Bor | Bor | -17.9 |
| Kürnberg | Ind | Ind | + 6.1 |
| Mühlhauser Halde | Ind | Ind | -6.7 |

However, when assigning the sites to the four dominance categories, depending on whether the highest mean biomass was found at the *arable land* (AL = MT1), the *border* between arable land and nature protected area (Bor = MT2), within *nature protected areas* (NPA = MT3–5), or *indistinguishable* with similarly high values for at least two categories (Ind = MT1–MT2–MT3–5), the annual repeatability was moderately high with 12 out of 21 sites showing similar dominance patterns for both sampling periods (Table 1).

While the categorial biomass distribution was stable between years for 12 out of 21 sites, there were changes at the other nine sites from 2020 to 2021 (Table 1). A bias in the data was caused by the withdrawal of the farmers’ consent, which resulted in the trap on the arable land (MT1) having to be abandoned at six sites and moved to the borders during data acquisition. At one site (Brauselay) MT1 had to be removed in 2021 completely. Interestingly, this had little impact on the biomass dominance distribution, as in five cases the pattern was identical to that of 2020 and only in one case the highest biomass was shifted from an indistinguishable distribution towards the border. Therefore, the bias due to the relocation of the trap from the arable land in the second year seems to be small. Overall, the insect biomass was higher only on arable land at a site with environmentally friendly farming and a species rich vegetation including many endangered plants (Köthe et al., 2023b; Table 1: Oderhänge Mallnow). In two cases (Riedensee, Insel Koos), the highest mean biomass was found in the second year 2021 on previous arable land, and not anymore in the NPA, which can be attributed to a shift from arable land to newly established wildflower strips. At some sites, flowered fallow land and uncut ‘flowering islands’ around Malaise traps, led to a shift in biomass distribution from one year to the next (particularly pronounced at Bottendorfer Hügel and Schwellenburg; Table 1 and Figure S1). At most sites, the highest biomass occurred equally in the NPA or at the border, or showed no clear pattern and was classified as indistinguishable (Ind) (Table 1, Supplementary Table S3).

Overall, the mean insect biomass per day in 2020 was highest at the most central point within a NPA along the transect (MT5) with 2.21 g/d, and the highest maximum biomass per day with 7.62 g/d (Figure 2 left). The situation was different in 2021. Due to newly established wildflower strips, flowered fallow land and uncut ‘flowering islands’ around four of the Malaise traps (MT1), the original experimental setup was distorted. The mean insect biomass was highest on arable land (MT1) at 2.05 g/d and the maximum biomass of 8.19 g/d was highest at the border (MT2; Figure 2 right).

**FIGURE 2.**
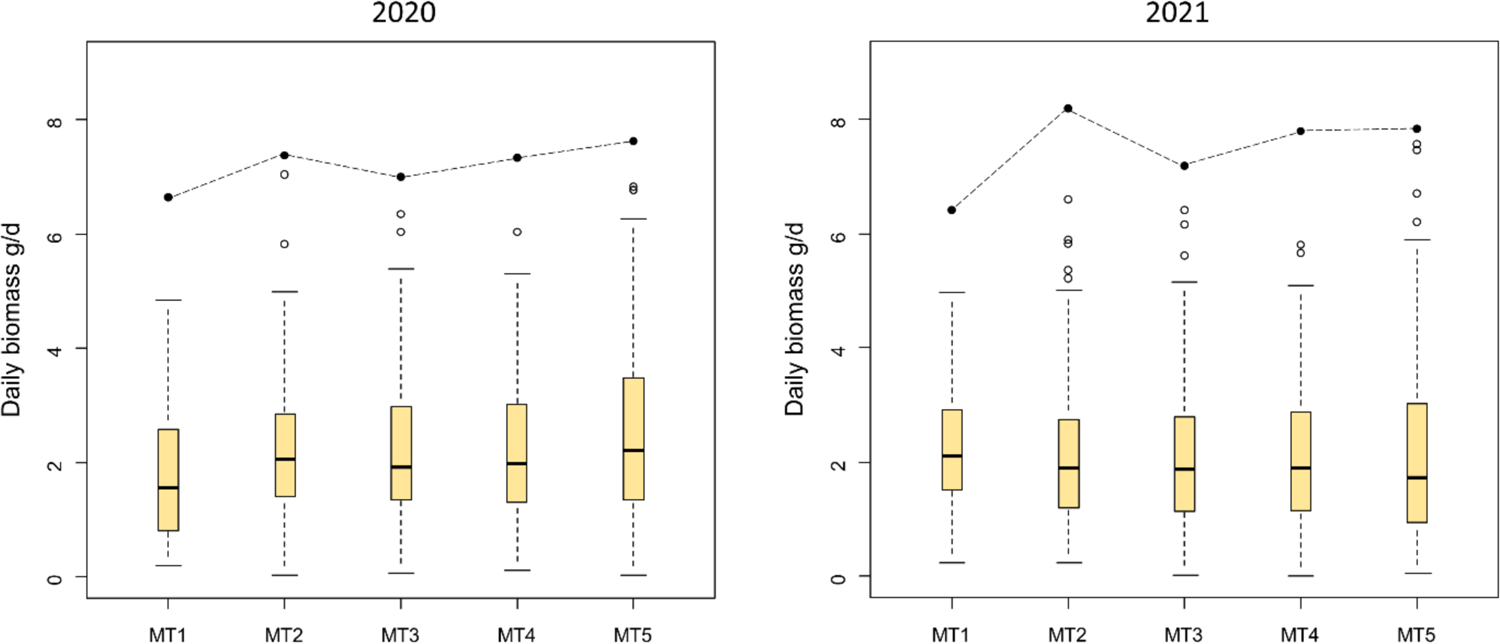
Boxplots of the median insect biomass along the Malaise trap transect for the sampling intervals 2020 (left) and 2021 (right). Full circles at the top represent the maximum biomass.

To enable a better comparison with previous similar studies in German NPA, we have limited the following analyses to the traps MT3 to MT5 (with a total of 500 samples from 2020 and 475 from 2021). These traps were located within the protected areas, while traps on borders and in adjacent arable land were excluded. Generalized additive models revealed that three indices of climate (mean maximum temperature, mean precipitation, and continentality index) were not associated with insect biomass (Figure 3A–D), regardless of the model. Agricultural production area within 2 km around NPA tended to be negatively associated with insect biomass in 2020 (P = 0.07; Figure 3A) but not in 2021 (P = 0.72; Figure 3B). Excluding the four sites with newly established ‘flower strips’ in 2021, the agricultural production area within a radius of 2 km had a no significant negative effect on the mean daily insect biomass in both years (2020: P = 0.10, Figure 3C; 2021: P = 0.32, Figure 3D).

**FIGURE 3.**
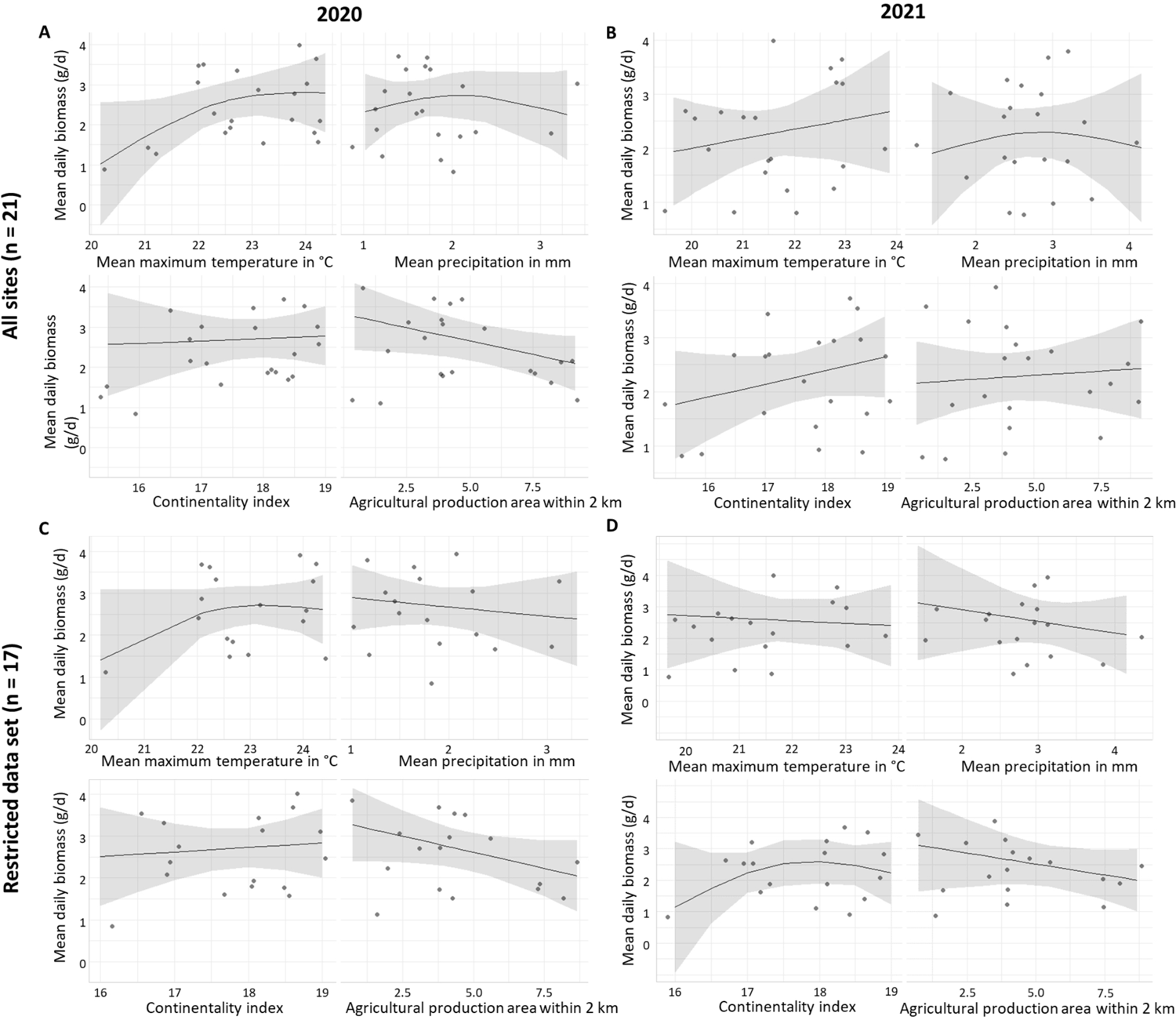
Results of generalized additive models (GAM) that examine the effects of mean maximum temperature, mean precipitation, continentality index and agricultural production area (2 km radius around NPA) on mean insect biomass per day within NPA (pooled data from MT3–5). Model A includes all 21 sites for 2020, and model B for 2021. Model C excludes four sites with newly established “flower strips” (Riedensee, Insel Koos, Bottendorfer Hügel, Schwellenburg) for 2020, and model D for 2021.

In DINA, the malaise traps were emptied 189 times across the 21 sites in 2020 and 165 times in 2021. The samples cover various biogeographical regions in Germany, including potentially more species-rich areas in southern Germany. Despite selecting high-quality NPA with endangered plant communities and biotope types according to the Red Lists for Germany, our measurements show a comparable low level of biomass according to the data from the earlier period (2013–2016; Hallmann et al. 2017; Figure 4).

**FIGURE 4.**
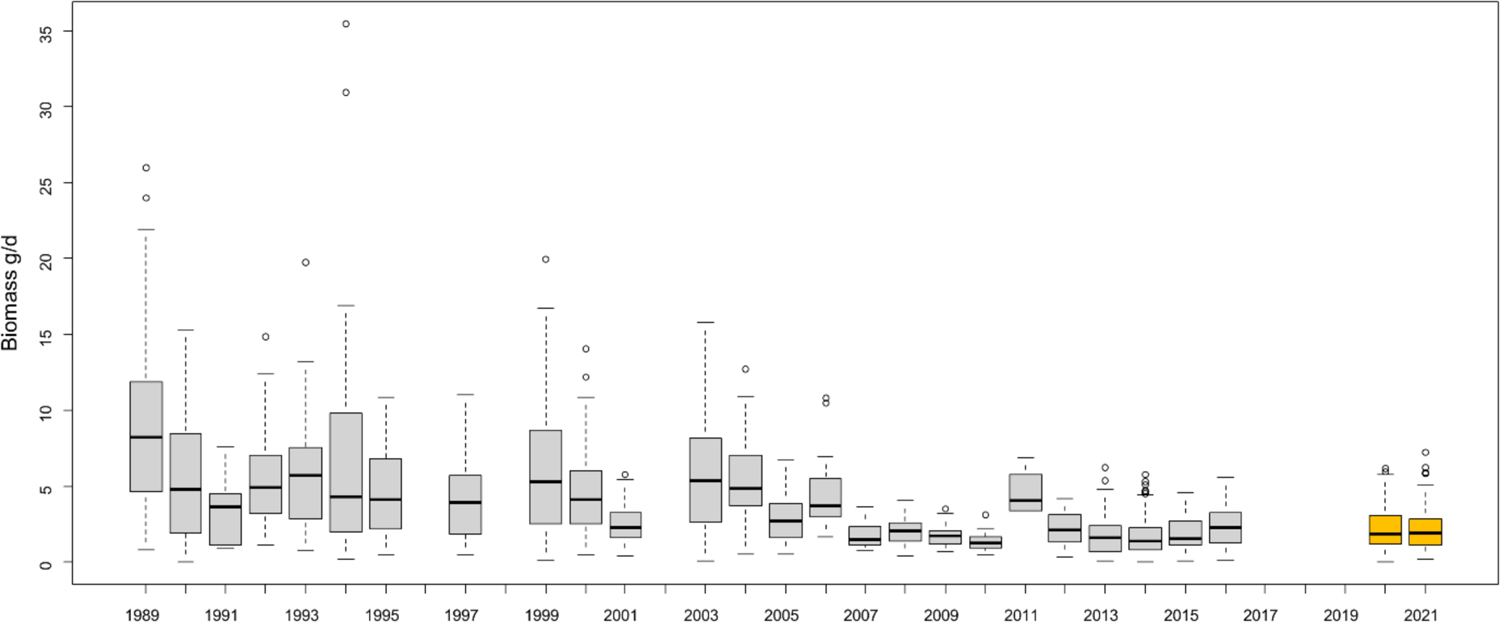
Daily insect biomasses in gram in German nature protected areas (NPA) of a 27-year timeseries published in Hallmann et al., (2017) (grey) compared to our DINA data (mean daily biomass of MT3–5) from 2020 and 2021 (orange).

## Discussion

Our study shows a relatively low biomass of flying insect in 21 nature protected areas spread across Germany. Insect biomass levels were similarly low as during the last years in the previous study by Hallmann et al. (2017) over a period of 27 years. While using the same standardised methods and traps, Hallmann et al. (2017) focused on NPA mainly in three German federal states, two in the west (North Rhine-Westphalia, Rhineland Palatinate) and one in the east (Brandenburg). In contrast, our DINA project selected NPA representatively across Germany (Lehmann et al., 2021) including some in warmer climates in southern Germany where insect species richness is generally higher. We did not find significant regional differences among the 21 sites. In line with another recent study using different types of Malaise traps (Welti et al., 2022), we could also observe that mean maximum temperatures correspond positively but not significantly with detected biomass. The differences between the two monitoring periods of 2020 and 2021 were considerably in some sites (Table 1, Supplementary Figure S2) but no general trend can be derived by our results. Fluctuations of insect populations are a well-known natural phenomenon and one of the major challenges in interpreting insect monitoring data (Didham et al., 2020).

The reasons for the deviations in biomass observed by us between the two consecutive years seem to be diverse, and related to local weather events (storms, heath waves, etc.) and anthropogenic influences. After the first year of the DINA project, some farmers changed their habits and started planting flower strips near our traps to support insects, changed crop management, or stopped agricultural cultivation (see Supplementary Table S1). We could clearly observe immediate consequences at two sites (Riedensee, Insel Koos), where flower strips were established in 2021 and possibly initiated as a response to DINA. In both cases, the maximum insect biomass was shifted from protected habitats towards the previously cultivated arable land (Table 1, Supplementary Figure S1) and can be explained by the change of local land use around this trap. A flowered fallow was created at the Bottendorfer Hügel site in 2021, while the vegetation around MT 1 and 2 was not cut at the Schwellenburg site in 2021, so that distinctive flowering islands have formed around the Malaise traps (Figure S3). In both cases, we could register shifting effects on insect biomass distribution along the transect. In the first year the maximum biomass was close to the centre of the NPA, while in the second year the maximum were at the edge or even outside the NPA. Flower stripes have a major impact on insect abundance, increasing the number of individuals and species (Jönsson et al., 2015; Lowe et al., 2021). However, these specimens are lured from their natural habitats into the nearby arable land, potentially increasing the risks of exposure to insecticides and other pesticides (Botías et al., 2016; Brühl et al., 2021; Hahn et al., 2015). These measures and effects were taken into account in our calculation. The model, which excluded the sites with such measures, showed an association between the proportion of arable land within a radius of 2 km around NPA. Different field management methods can also influence the occurrence of insects. Maize, for example, tends to result in higher arthropod biomass compared to other cultivations (Frizzas et al., 2018; Hüber et al., 2022; Musters et al., 2021; Sorribas et al., 2016). Almost no changes in insect biomass (–1%; Table 1, Supplementary Figure S1) were recorded at the Wipperdurchbruch site between 2020 and 2021, regardless of land use changes and crop rotation, probably due to the fact that the NPA has no direct contact with arable land.

Although higher biomass prevailed in the nature protected areas, one third of our sites had the maximum measured biomass at the borders. This can be related to edge effects, as many flying insects patrol along prominent terrain boundaries. Such boundaries can be the break between relatively tall cereals and low natural vegetation, as is often seen at field edges. But even differences that are not visible to humans, such as differences in plant species compositions, can be perceived by insects as borderlines (Macfadyen & Muller, 2013; Nguyen & Nansen, 2018). Because insects stay near chemically treated fields for longer periods, they face a higher risk to be contaminated with insecticides and other pesticides. Few sampling sites had the maximum insect biomass on arable land. At the Oderhänge Mallnow site, the environmentally friendly farming practises over years led to a high diversity of plants including endangered species typical for arable land (Köthe et al., 2023b).

On average, we observed an increase in maximum insect biomass per day along the transect from the edge towards the centre of the nature protected area in 2020 and from MT3 to MT4 in 2021. These findings emphasise our previous results on spillover effects into the NPA (Köthe et al., 2023b), which documented strong chemical edge effects and negative impacts from adjacent arable land on plant communities in nature protected areas. In addition, the number of endangered plant species rose with increasing proximity to the edge of the field. This strong influence of intensive cultivation around and in nature protected areas, both for insects and endangered plant species, calls for more effective buffer zones around NPA.

## Conclusions

Our study confirms the low level of insect biomass in German nature protected areas compared to available data two to three decades ago. Furthermore, our results show that nature reserves act as important habitats for insects with maximum biomasses closer to the centre of these areas. Nevertheless, they are often not sufficiently buffered against negative influences from directly adjacent, conventionally farmed croplands. The indication of negative lateral effects corresponds to a higher number of endangered plant species and a decrease in nitrogen indicator values along the transects. Appropriate measures must therefore be taken to better implement the conservation objectives of nature reserves and their habitats, as these areas are often the last refuges of critically endangered species.

Changes in land use such as newly established flower strips and bee friendly cultivations as measures with the aim to improve the situation for the insects, had an immediate impact in our study. Increased edge effects are seen at the immediate borders of cultivated areas. Many insects patrol along these and this behavioural pattern of highly active species brings large numbers of insects close to pollution sources, which contradicts nature conservation goals. Therefore, such measures must be planned and linked to other accompanying actions in order to provide the insects with the best possible food and reproduction bases and to protect them from exposure to pesticides.

The decline in insect biomass in German nature reserves shows that insects, which perform key functions in our ecosystems, need better protection and that efforts so far have not been able to reverse the loss.

## Acknowledgments

The Project DINA is funded by the German Federal Ministry Education and Research (BMBF) and is handled by the VDI Project Management Agency (grant number FKZ 01LC1901). Conceptual framework and development of methodologies of the Entomological Society Krefeld (EVK) was funded by the German Federal Ministry for the Environment, Nature Conservation, Nuclear Safety and Consumer Protection (BMUV), handled by the Bundesamt für Naturschutz (BfN), grant number FKZ 3516850400. We are grateful to farmers and authorities for granting permits to install Malaise traps and to take samples. Finally, the project would not have been possible without the dedicated support of our over 50 volunteers and Citizen Scientists who maintained the Malaise traps.

## Author contributions

The study was conceptualized by GUCL and MS. RM, SK and GUCL wrote the original draft. Development and design of methodology as well as data collection was conducted by all co-authors. LE prepared the climatic spatial data. SK analysed the data. All co-authors reviewed and edited the document and approved the final manuscript.

## Data availability

The data generated and/or analysed as well as R codes can be made available for research purposes upon request to the corresponding authors.

## Declarations

Conflict of interest: The authors declare that they have no conflict of interest.

## Supplement

**FIGURE S1.**
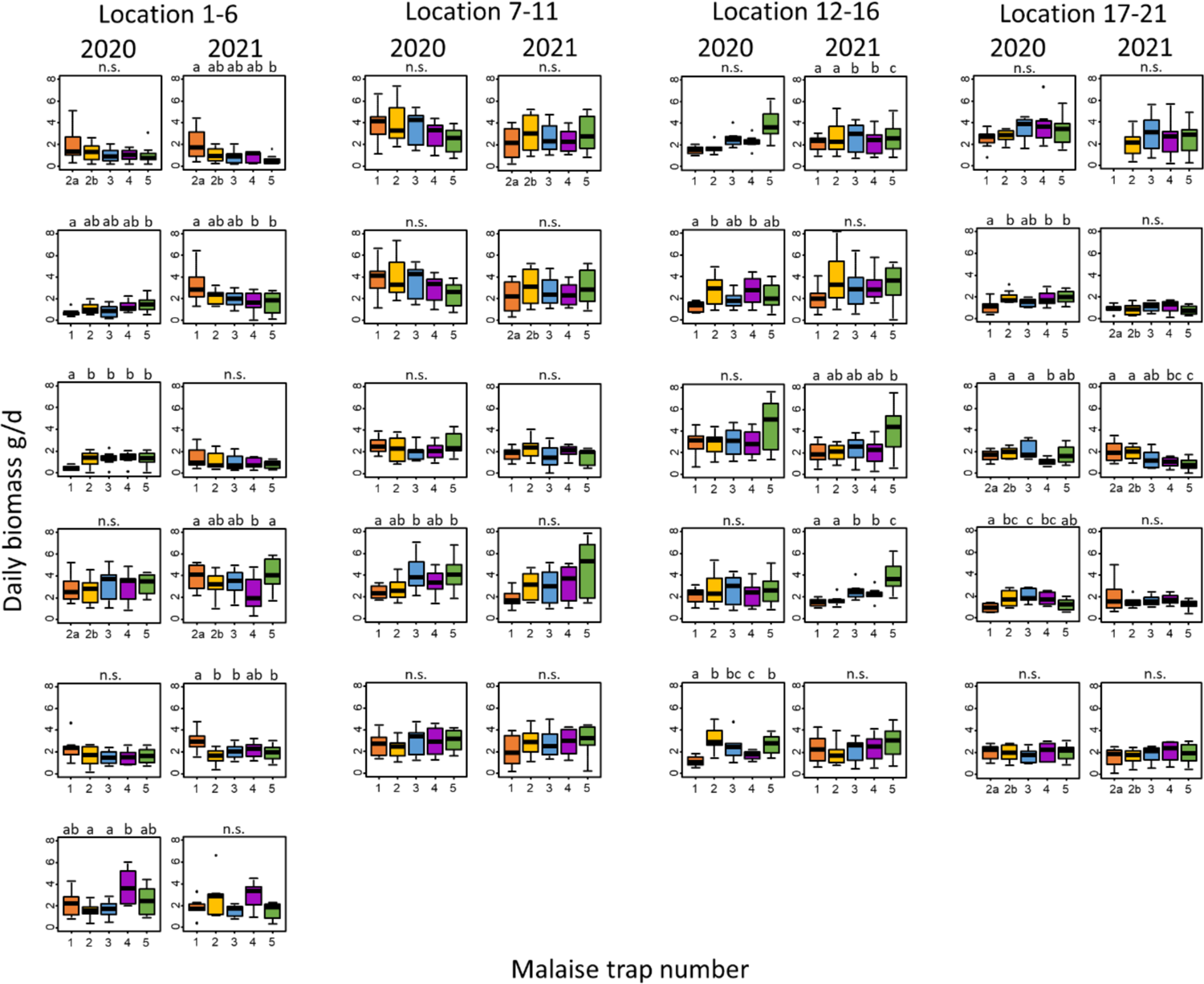
Mean daily biomass per trap at each sampling site for 2020 and 2021. Lower case letters classify differing groups with a significance level of p < 0.05 according to post-hoc Mann-Whitney U-tests. Location numbers according to Table S2.

**FIGURE S2.**
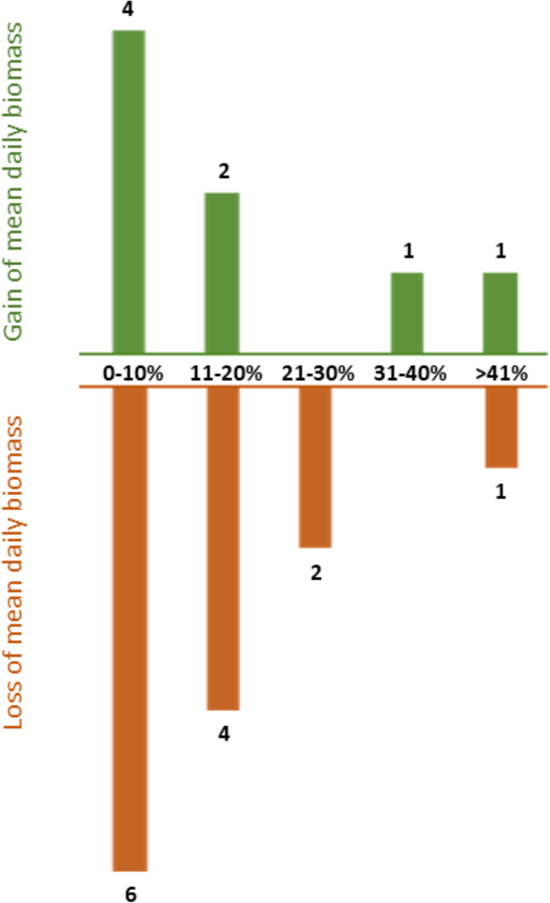
Number of locations with gain (green) or loss (orange) of daily biomass in percent.

**FIGURE S3.**
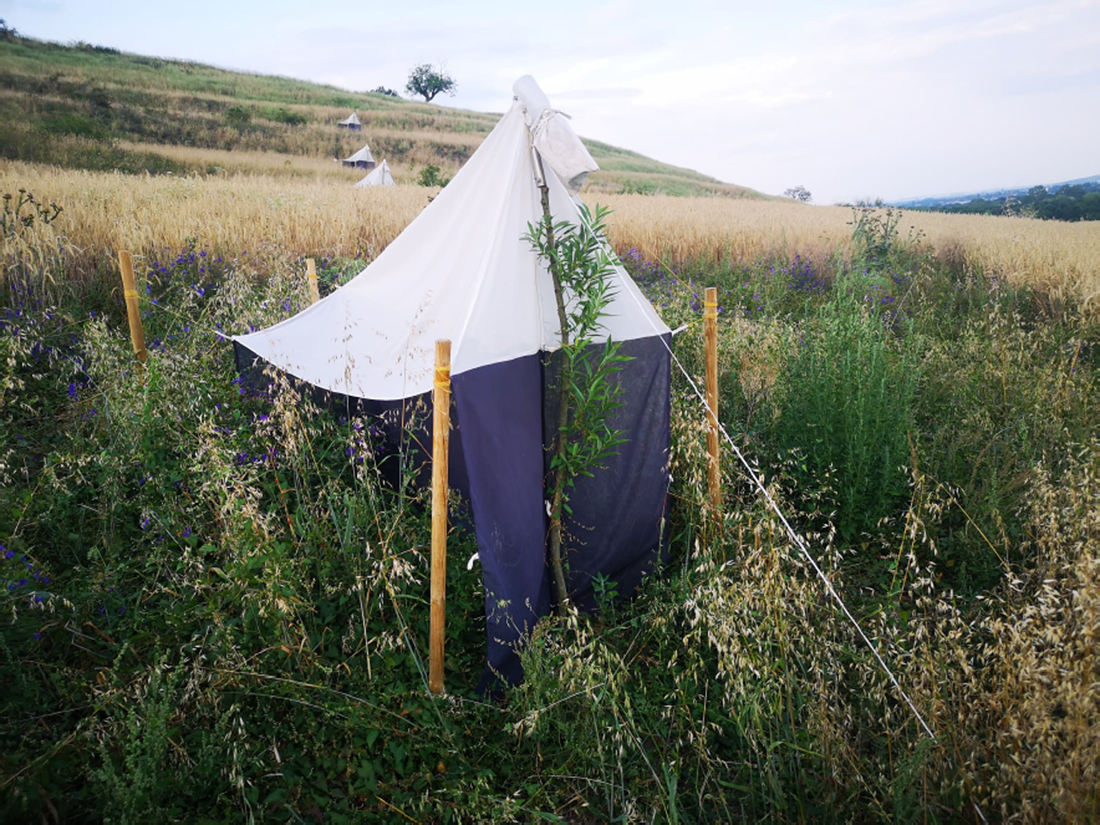
Cereal field with island of wildflowers around Malaise trap 1 (MT1) (Schwellenburg, Thuringia; 26.07.2021)

---

**TABLE S1.** Mean daily biomass with standard deviation and sample size of the two sampling years 2020 and 2021, the percentual difference between the two years.

| Location | Mean daily biomass 2020 | Mean daily biomass 2021 | Difference (in %) | Site-specific comments | Further comments (trap operations) |
| --- | --- | --- | --- | --- | --- |
| 01_LUE | 1.25 ± 0.99<br>(n = 35) | 1.14 ± 0.97<br>(n = 40) | -8.8 | Tipped transect in both years | No special observations. Minor damages. |
| 02_RIE | 1.09 ± 0.60<br>(n = 39) | 2.12 ± 1.14<br>(n = 40) | +51.4 | Storm damages | No special observations. Team of same three people throughout both seasons. |
| 03_KOO | 1.13 ± 0.66<br>(n = 39) | 1.10 ± 0.76<br>(n = 35) | -2.7 | 2021 flower strip at MT1 | Every year with new team. Many damages in spring 2020 due to cattle. Numerous wind damages throughout both seasons. |
| 04_GEE | 3.00 ± 1.23<br>(n = 38) | 3.38 ± 1.41<br>(n = 40) | +11.2 | Tipped transect in both years | No specials observations. Minor damages. |
| 05_MAL | 1.72 ± 0.85<br>(n = 40) | 2.15 ± 0.86<br>(n= 40) | +20.0 |  | No specials observations. Minor damages. |
| 06_WIS | 2.35 ± 1.36<br>(n = 40) | 2.10 ± 1.31<br>(n = 35) | -10.6 |  | Change of trap operator in 2020. Occasionally supported by 07_BIS team. |
| 07_BIS | 3.45 ± 1.43<br>(n = 40) | 3.62 ± 1.98<br>(n = 28) | +4.7 | Flooding in 2021 | Team of same people throughout both seasons. |
| 08_GIP | 3.31 ± 1.58<br>(n = 39) | 2.67 ± 1.41<br>(n = 40) | -19.3 | Tipped transect in 2021 | No special observations. Frequently problems with farmer at MT1. |
| 09_POR | 2.31 ± 0.92<br>(n = 39) | 1.85 ± 0.86<br>(n = 36) | -19.9 |  | Numerous storm damages throughout both seasons. Special fixations of traps (bamboo sticks) due to very hard soil. |
| 10_ZIE | 3.34 ± 1.38<br>(n = 40) | 3.19 ± 1.87<br>(n = 38) | -4.5 |  | No special observations. Minor damages. |
| 11_WIP | 2.88 ± 1.45<br>(n = 40) | 2.85 ± 1.66<br>(n = 39) | -1.0 |  | No special observations. Minor damages. |
| 12_BOT | 1.68 ± 0.98<br>(n = 38) | 1.56 ± 0.68<br>(n = 38) | -7.1 | 2021 no agricultural practice | Numerous wind damages throughout both seasons. |
| 13_SBG | 2.14 ± 1.18<br>(n = 40) | 3.06 ± 1.84<br>(n = 38) | +30.1 |  | Area around MT1 & MT2 not cultivated. Many flowers around traps in the second year. |
| 14_HOF | 3.24 ± 1.61<br>(n = 39) | 2.50 ± 1.54<br>(n = 40) | -22.8 |  | Change of trap operator team in spring 2021. |
| 15_KOP | 2.37 ± 1.14<br>(n = 40) | 2.48 ± 1.22<br>(n = 40) | +4.4 |  | No special observations. |
| 16_DOE | 2.21 ± 1.11<br>(n = 40) | 2.33 ± 1.22<br>(n = 40) | +5.1 |  | Change of trap operator team in spring 2021. |
| 17_BRA | 3.16 ± 1.21<br>(n = 40) | 2.49 ± 1.54<br>(n = 32) | -21.2 | MT1 missed in 2021 | Very difficult terrain. No special observations. |
| 18_MIT | 1.65 ± 0.65<br>(n = 40) | 0.95 ± 0.47<br>(n = 39) | -42.4 | Tipped transect in 2021 | Conflict with farmer in summer 2020: dislocation of MT1 |
| 19_IPF | 1.73 ± 0.69<br>(n = 39) | 1.42 ± 0.83<br>(n = 40) | -17.9 | Tipped transect in both years | Change of trap operator in spring 2021. |
| 20_KUE | 1.55 ± 0.70<br>(n = 40) | 1.65 ± 0.78<br>(n = 40) | +6.1 |  | No special observations. |
| 21_MUE | 1.95 ± 0.70<br>(n = 39) | 1.82 ± 0.82<br>(n = 40) | -6.7 | Tipped transect in both years | No special observations. |

**TABLE S2.** Location information.

| Nr | Abbrev. | Location | State | FFH (SAC) -ID *1 | FFH (SAC) Area [ha] *1 | NSG-ID *2 | NSG Area [ha] *2 | Protected habitat types (the transect is crossing) *3 | Endangered plant communities along transect (German Red List with status) *4 |
| --- | --- | --- | --- | --- | --- | --- | --- | --- | --- |
| 1 | LUE | Lütjenholmer Heidedünen | SH | 1320-302 | 313 | 37570 | 18 | Dry sand heaths with Calluna and Genista, Northern Atlantic wet heaths with Erica tetralix | Ericetum tetralicis (2); Myricetum gale (2); Genisto pilosae-Callunetum (2) |
| 2 | RIE | Riedensee | MV | 1836-301 | 113 | 33182 | 120 | Xeric sand calcareous grasslands |  |
| 3 | KOO | Insel Koos | MV | 1747-301 | 60406 | 33165 | 1574 | Lowland hay meadows (Corresponding categories: German classification BTT-Nr. 34070102 species-rich, fresh pasture of the planar to submontane level) | Festuco rubrae-Cynosuretum cristati (3); |
| 4 | GEE | Geesower Hügel | BB | 2752-301 | 82 | 30009 | 39 | Xeric sand calcareous grasslands, Sub-pannonic steppic grasslands | Potentillo arenariae-Stipetum capillatae (2) |
| 5 | MAL | Oderhänge Mallnow | BB | 3552-306 | 305 | 30118 | 305 | Xeric sand calcareous grasslands, Sub-pannonic steppic grasslands | Adonido vernalis-Brachypodietum pinnati (2) |
| 6 | WIS | Wisseler Dünen | NW | 4203-301 | 71 | 35714 | 79 | Inland dunes with open Corynephorus and Agrostis grasslands | Airetum praecocis (3) |
| 7 | BIS | Bislicher Insel | NW | 4305-301 | 1002 | 36965 | 1053 | Rivers with muddy banks with Bidention-Vegetation, Hydrophilous tall herb fringe communities of plains |  |
| 8 | GIP | Gipskarstlandschaft Hainholz | NI | 4226-301 | 1327 | 33265 | 641 | Lowland hay meadows | Festuca rubra-Agristis capillaris (3) |
| 9 | POR | Porphyrlandschaft bei Gimritz | ST | 4437-302 | 819 | 38460 | 288 | Sub-pannonic steppic grasslands, Siliceous rock with pioneer vegetation | Festuco valesiacae-Stipetum capillatae (2) |
| 10 | ZIE | Ziegenbuschhänge bei Oberau | SN | 4847-301 | 112 | 37981 | 20 | Lowland hay meadows | Arrhenateretum elatioris (V) |
| 11 | WIP | Wipperdurchbruch | TH | 4631-302 | 6869 | 38140 | 672 | Semi-natural (semi-)dry grasslands and scrubland facies on calcareous substrates (Festuco-Brometalia) | Xerobrometum (3) |
| 12 | BOT | Bottendorfer Hügel | TH | 4634-303 | 133 | 38143 | 134 | Calaminarian grasslands of the Violetalia calaminariae, Sub-pannonic steppic grasslands | Armerietum bottendorfensis (3) |
| 13 | SGB | Schwellenburg | TH | 4931-301 | 89 | 38111 | 23 | Rupicolous calcareous or basiphilic grassland of the Alyso-Sedion albi, Sub-pannonic steppic grasslands | Brometum (2) |
| 14 | HOF | Hofberg | TH | 5327-305 | 260 | 38319 | 43 | Rupicolous calcareous or basiphilic grassland of the Alyso-Sedion albi, Semi-natural dry grasslands and scrubland facies on calcareous substrates (Festuco-Brometalia) | Arrhenateretum elatioris (V); Brometum (2); Gentiano-Koelerietum pyramidatae (3) |
| 15 | KOP | Koppelstein - Helmestel | RP | 5711-301 | 4551 | 37188 | 87 | Semi-natural (semi-)dry grasslands and scrubland facies on calcareous substrates (Festuco-Brometalia), Lowland hay meadows | Arrhenateretum elatioris (V); Brometum (2) |
| 16 | DOE | Rheinhänge Dorscheider Heide | RP | 5711-301 | 4551 | 37186 | 626 | European dry heaths, Siliceous rock with pioneer vegetation of the Sedo-Scleranthion | Airo caryophylleae-Festucetum ovinae (3); Thymus-Festuca (3) |
| 17 | BRA | Brauselay | RP | 5809-301 | 16273 | 37136 | 14 | Stable xerothermophilous formations with Buxus sempervirens on rock slopes (Berberidion), Silicate rocks with pioneer grassland, Tilio-Acerion forests of slopes, screes and ravines |  |
| 18 | MIT | Mittelberg | BW | 7319-341 | 853 | 30714 | 45 | Semi-natural (semi-)dry grasslands and scrubland facies on calcareous substrates (Festuco-Brometalia) | Brometum (2) |
| 19 | IPF | Ipf | BW | 7327-341 | 3363 | 30585 | 60 | Juniperus communis formations on calcareous grasslands, Semi-natural dry grasslands and scrubland facies on calcareous substrates (Festuco-Brometalia) | Arrhenateretum elatioris (V); Brometum (2) |
| **20** | **KUE** | **Kürnberg** | BW | 7427-341 | 990 | 30689 | 12 | Juniperus communis formations on calcareous grasslands, Semi-natural (semi-)dry grasslands and scrubland facies on calcareous substrates (Festuco-Brometalia) | Festuca rubra-Agrostis capillaris (3); Brometum (2) |
| **21** | **MUE** | **Mühlhauser Halde** | BW | 7916-311 | 3678 | 31149 | 52 | Juniperus communis formations on calcareous grasslands, Semi-natural (semi-)dry grasslands and scrubland facies on calcareous substrates (Festuco-Brometalia) | Arrhenatheretum elatioris (V); Brometum (2) |

\*1 https://www.bfn.de/themen/natura-2000/natura-2000-gebiete/steckbriefe.html#c33722
\*2 https://geodienste.bfn.de/schutzgebiete?lang=de&layers=NSG
\*3 EU Commission (2013): EUR 28 Interpretation Manual of European Union habitats
\*4 Rennwald, E. et al. (2002). Rote Liste der Pflanzengesellschaften Deutschlands mit Anmerkungen zur Gefährdung. Schriftenreihe für Vegetationskunde. 35. 393-592.

**TABLE S3.** Distribution of categories I - IV according to the maximum mean biomass. **AL** = arable land, **Bor** = boarder of arable land and nature reserve, **NPA** = within nature protected area, **Ind** = indistinguishable (at least two categories show similarly high values).

| Category | 2020 | 2021 | Total |
| --- | --- | --- | --- |
| AL | 1 | 3 | 4 |
| Bor | 6 | 7 | 13 |
| NPA | 8 | 3 | 11 |
| Ind | 6 | 8 | 14 |
|  | 21 | 21 | 42 |

**TABLE S4.** Values of the Generalized additive models (GAM) from Figure 3.

| Model A | Variable | Edf | F value | P value |
| --- | --- | --- | --- | --- |
|  | Temperature | 1.64 | 1.33 | 0.22 |
|  | Precipitation | 1.47 | 0.54 | 0.64 |
|  | Continentality index | 1.00 | 0.06 | 0.81 |
|  | Agricultural production area in 2 km | 1.00 | 3.81 | 0.07 |
| Model B | Temperature | 1.00 | 0.56 | 0.47 |
|  | Precipitation | 1.36 | 0.27 | 0.76 |
|  | Continentality index | 1.00 | 1.34 | 0.27 |
|  | Agricultural production area in 2 km | 1.00 | 0.13 | 0.72 |
| Model C | Temperature | 1.58 | 0.69 | 0.40 |
|  | Precipitation | 1.00 | 0.35 | 0.57 |
|  | Continentality index | 1.00 | 0.12 | 0.73 |
|  | Agricultural production area in 2 km | 1.00 | 2.65 | 0.13 |
| Model D | Temperature | 1.00 | 0.06 | 0.81 |
|  | Precipitation | 1.00 | 0.52 | 0.49 |
|  | Continentality index | 1.50 | 0.78 | 0.48 |
|  | Agricultural production area in 2 km | 1.00 | 1.08 | 0.32 |

